# Continuous directed evolution of a feedback-resistant *Arabidopsis* arogenate dehydratase in plantized *E. coli*

**DOI:** 10.1101/2022.09.24.509247

**Authors:** Bryan J. Leong, Andrew D. Hanson

## Abstract

Continuous directed evolution (CDE) is a powerful tool for enzyme engineering due to the depth and scale of evolutionary search that it enables. If suitably controlled and calibrated, CDE could be widely applied in plant breeding and biotechnology to improve plant enzymes *ex planta*. We tested this concept by evolving *Arabidopsis* arogenate dehydratase (AtADT2) for resistance to feedback inhibition. We used an *Escherichia coli* platform with a phenylalanine biosynthesis pathway reconfigured (‘plantized’) to mimic the plant pathway, a T7RNA polymerase-base deaminase hypermutation system (eMutaT7), and 4-fluorophenylalanine as selective agent. Selection schemes were pre-validated using a known feedback-resistant AtADT2 variant. We obtained variants that had 4-fluorophenylalanine resistance at least matching the known variant and that carried mutations in the ACT domain responsible for feedback inhibition. We conclude that *ex planta* CDE of plant enzymes in a microbial platform is a viable way to tailor characteristics that involve interaction with small molecules.

## INTRODUCTION

Classical directed evolution has greatly advanced protein engineering^1,2^ but is limited by dependence on labor-intensive cycles of *in vitro* generation of sequence diversity and screening of variants.^3,4^ Classical directed evolution campaigns can thus only handle relatively few evolutionary cycles and independent evolving populations, and tend to incrementally climb local adaptive peaks rather than explore the fitness landscape more widely because they lack the mutational power needed to cross fitness valleys.^3–5^ Continuous directed evolution (CDE) systems overcome these limitations of depth and scale by hypermutating the target gene *in vivo* and, for enzymes, by coupling activity of the target gene to growth of the platform cell.^3,4^ CDE systems thus require only serial subculturing because they work by selecting for growth alone, and in principle they can run indefinitely.^3–5^ So far, CDE systems are largely confined to *Escherichia coli* and yeast,^4^ and while progress in CDE in mammalian cells^6^ and plants^7^ continues, long generation times and small library sizes are enduring obstacles.

Tailoring enzyme properties by CDE has many potential uses in plant breeding and plant biotechnology, including developing new herbicide-resistance genes, extending enzyme life, and relieving feed-back inhibition of flux-determining enzymes.^7–10^ As CDE in a plant platform is far less powerful than in a microbial one, a preferable strategy is to run CDE on a plant enzyme in *E. coli* or yeast, then return the improved gene to the plant by genome editing.^9^ This strategy is still at the concept stage. Further, for the *E. coli* systems, the differences between conditions in plant and prokaryote cells (e.g., metabolic pathway architecture, metabolite levels, redox poise, protein-folding and degradation systems^8^) and the uncertain durability of the hypermutation machinery in long evolution campaigns^4,5,8^ are potential roadblocks. Nor is it clear that *E. coli* CDE systems work in minimal media, which is critical because many selection schemes require auxotrophs.^5^

We therefore undertook a proof-of-concept study in *E. coli* to evolve a plant arogenate dehydratase (ADT) (EC 4.2.1.91) for resistance to feedback inhibition by phenylalanine. We used eMutaT7,^11^ one of several CDE systems whose mutagenesis machinery is a T7RNA polymerase (T7RNAP)-nucleobase deaminase fusion that is directed to the target gene by a T7 promoter.^4^ The eMutaT7 system uses cytidine deaminase (CD), which makes mainly C→T mutations (**Figure 1A**). ADT was chosen as the target because (i) it requires rewiring (‘plantizing’) of the *E. coli* aromatic amino acid synthesis network to reproduce that in plants (**Figure 1B**), making the study more demanding and informative, and (ii) a known feedback-resistant ADT variant^12^ is available to pre-validate and calibrate selection schemes. Having confirmed that eMutaT7 works in minimal medium, we showed that short evolution campaigns deliver feedback-resistant ADT variants. This opens the way to custom applications of *ex planta* CDE using eMutaT7 and related systems to evolve plant enzymes to meet various needs.

**Figure 1.**
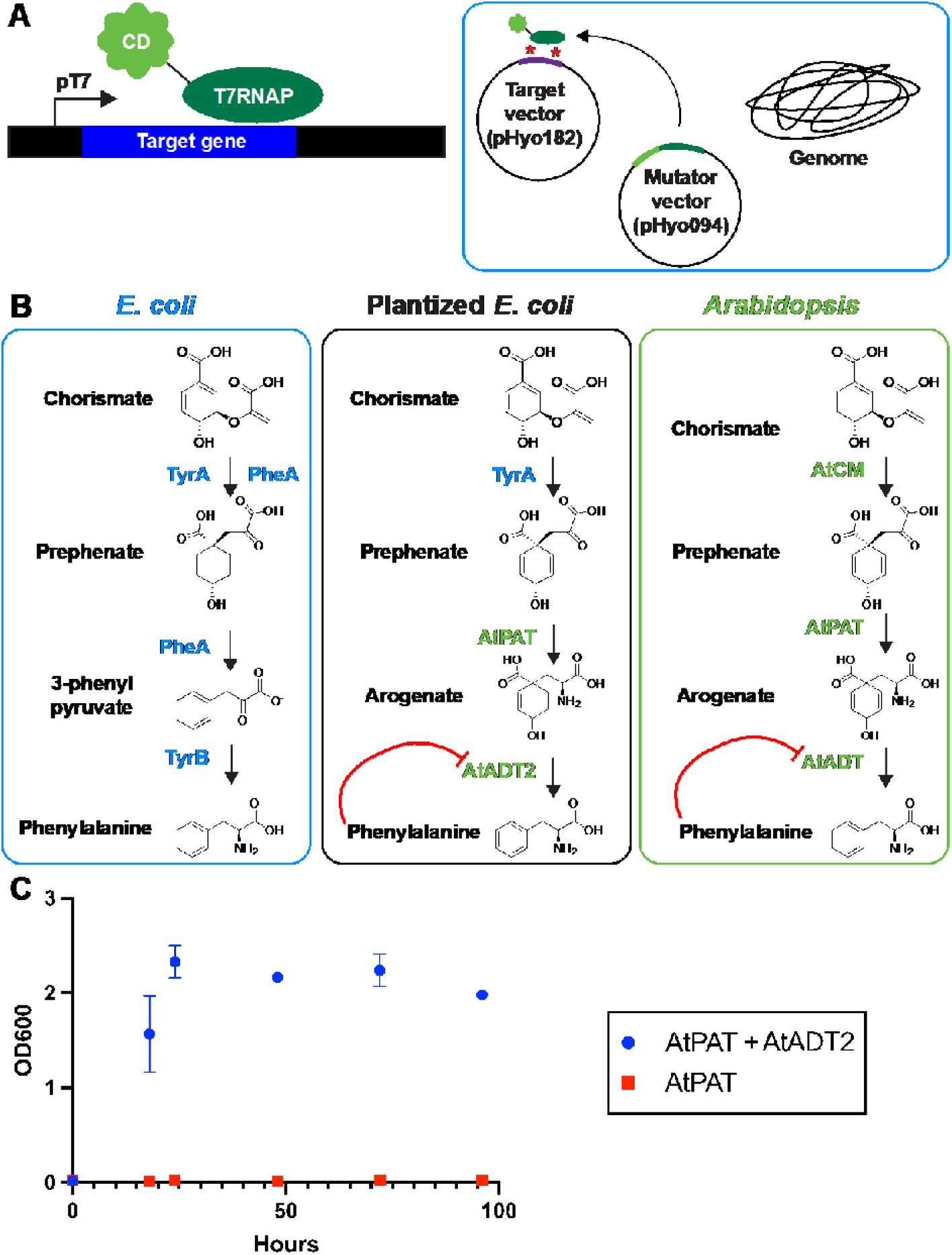
The eMutaT7 continuous directed evolution system and the plantized *E. coli* platform strain. (A) eMutaT7 hypermutates a plasmid-borne target gene using a T7RNAP-cytidine deaminase (CD) fusion that is encoded by a second plasmid. (B) The native *E. coli* (left), *Arabidopsis* (right), and plantized *E. coli* (center) phenylalanine synthesis pathways. The native pathways proceed from prephenate via 3-phenylpyruvate in *E. coli* and via arogenate in *Arabidopsis*. PheA, bifunctional chorismate mutase/prephenate dehydratase; TyrB, tyrosine aminotransferase; TyrA, chorismate mutase activity of TyrA; AtPAT, *Arabidopsis* prephenate aminotransferase; AtADT2, *Arabidopsis* arogenate dehydratase 2. (C) Growth assays in MOPS minimal medium comparing *E. coli* Δ*pheA* strains expressing AtPAT alone or with AtADT2. Data are means ± s.e.m. of three independent replicates.

## RESULTS AND DISCUSSION

### Function in minimal medium

As eMutaT7 has so far been used only in rich media^4,11^ and might be metabolically burdensome enough to prevent growth in minimal media,^5^ we first tested its operation in MOPS minimal medium. A Δ*ung* strain was used as platform; the target was the gene encoding PheS A294G, a tRNA ligase that sensitizes *E. coli* to *p*-chlorophenylalanine. A campaign of three serial passages in minimal medium (without selection) gave multiple *p*-chlorophenylalanine-resistant mutants that all contained C→T nonsense mutations in the *pheS* coding sequence (**Table S1**). This result confirms that eMutaT7 allows *E. coli* growth and efficient mutagenesis in minimal medium.

### Plantizing the *E. coli* platform

*E. coli* and plants such as *Arabidopsis* synthesize phenylalanine from prephenate, but by different routes (**Figure 1B**).^13^ *E. coli* converts prephenate to phenylalanine via 3-phenylpyruvate using the prephenate dehydrogenase activity of PheA and the aminotransferase TyrB, whereas *Arabidopsis* does so via arogenate using prephenate aminotransferase (AtPAT) and arogenate dehydratase (AtADT2), which phenylalanine feedback-inhibits.^13^ To plantize the *E. coli* pathway, we deleted *pheA* and inserted an *AtPAT* cDNA flanked by the constitutive PJ23101 promoter/B0032 RBS and the *rrnB* T1 terminator into the *flu l*ocus. The *AtADT2* gene, expressed from the eMutaT7 pHyo182 vector, completes the plantized pathway. The chorismate mutase activity of *E. coli* TyrA suffices to replace that of PheA in this pathway. Growth tests on MOPS minimal medium confirmed operation of the plantized pathway and its dependence on AtADT2 (**Figure 1C**).

### Pre-validating the selection scheme

We adopted a classical analog selection scheme^14^ using 4-fluorophenylalanine (4FPA), which – like phenylalanine – inhibits plant ADT activity^15^ and also inhibits *E. coli* growth.^16^ AtADT resistance to inhibition by 4FPA is predicted to lead to an expanded free phenylalanine pool that competes out 4FPA’s inhibitory effect. To confirm that this is so, we compared growth of the plantized strain expressing wildtype AtADT2 or its feedback-resistant mutant.^12^ In line with prediction, growth of the wildtype AtADT2 strain was blocked by a high concentration (5 mM) of 4FPA (**Figure 2A**), whereas the feedback-resistant AtADT2 strain grew well after a lag (**Figure 2B**). As 4FPA concentration was reduced below 5 mM, the growth difference between the strains diminished progressively (**Figure S1**). These results validate the selection scheme. As recommended,^4^ we also ran a pilot test to check that the selection scheme does not yield frequent cheater mutations, e.g., ones not in the target ORF. To this end, we modified the mutator vector to express T7RNAP alone instead of the T7RNAP-CD fusion, and compared the frequency of 4FPA-resistant mutant recovery from cultures harboring each vector. The T7RNAP-CD strain gave resistant colonies at a frequency of 6.0 × 10^−7^ whereas the T7RNAP strain frequency was only 2.6 × 10^−8^ (**Table S2**). This 23-fold difference further validates the selection scheme.

**Figure 2.**
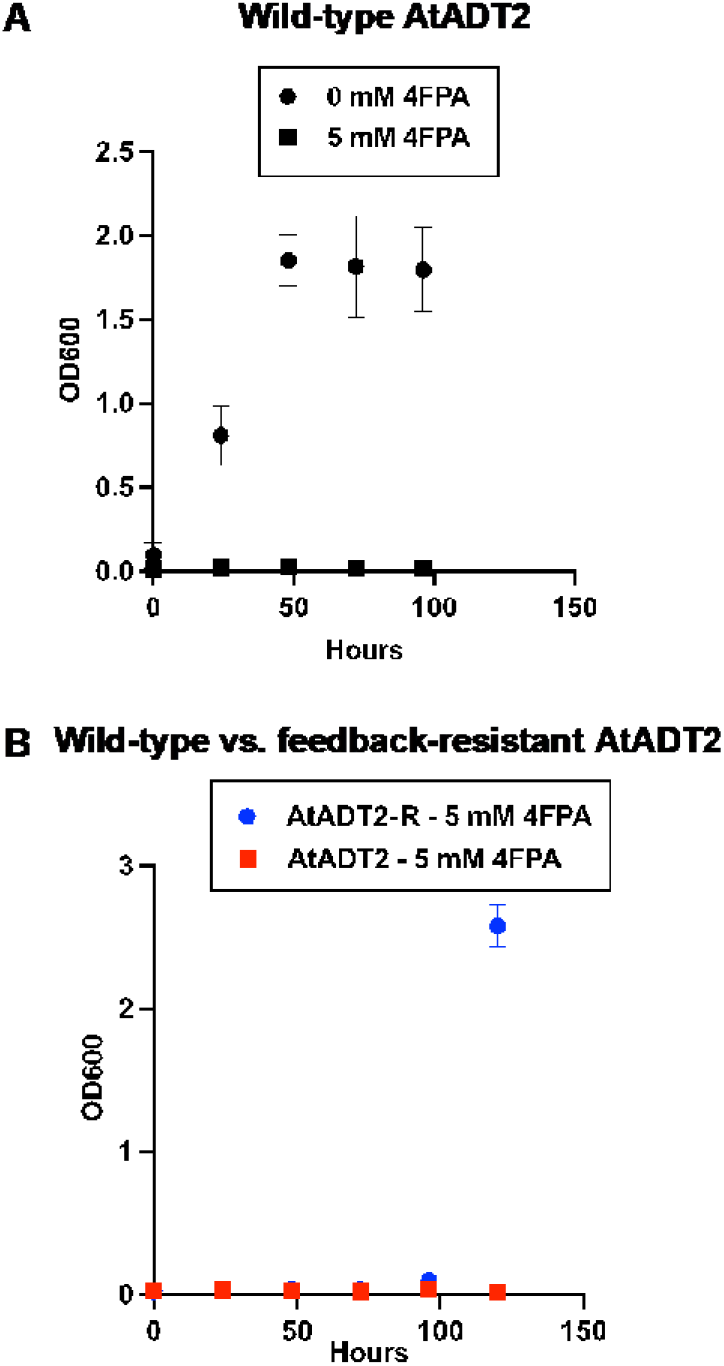
Use of the known feedback-resistant AtADT2 variant to pre-validate the planned select-ion scheme. (A) 4FPA-sensitivity of the plantized eMutaT7 strain expressing wildtype AtADT2. (B) Reduced 4FPA sensitivity of this strain expressing the feedback-resistant variant (AtADT2-R). Data are means ± s.e.m. of three independent replicates.

### Pilot eMutaT7 trial

We first ran eMutaT7 in the non-continuous mode used to develop it^11^ to confirm that feedback-resistant mutants could be recovered by building a mutant library and then screening it (**Figure 3**). Triplicate populations were mutagenized during three serial passages on non-selective LB medium, after which they were plated on selective MOPS minimal medium containing 5 mM 4FPA. Nine resistant colonies were isolated, restreaked on selective medium to eliminate residual non-resistant cells, and the AtADT2 ORFs were sequenced. Each ORF had one or other of six single or double mutations (**Figure 3 and Table S3**). Multiple mutants carried T190I or R213W mutations in AtADT2’s ACT feedback-regulation domain (**Figure 3**) in which the known feedback-insensitivity mutation is located.^12,17^ All mutations were C→T transitions (**Table S3**), as expected.^4^ In only one case was there a mutation in the promoter controlling AtADT2 expression. This trial confirmed that, in our application, the eMutaT7 mutational machinery performed as reported^12^.

**Figure 3.**
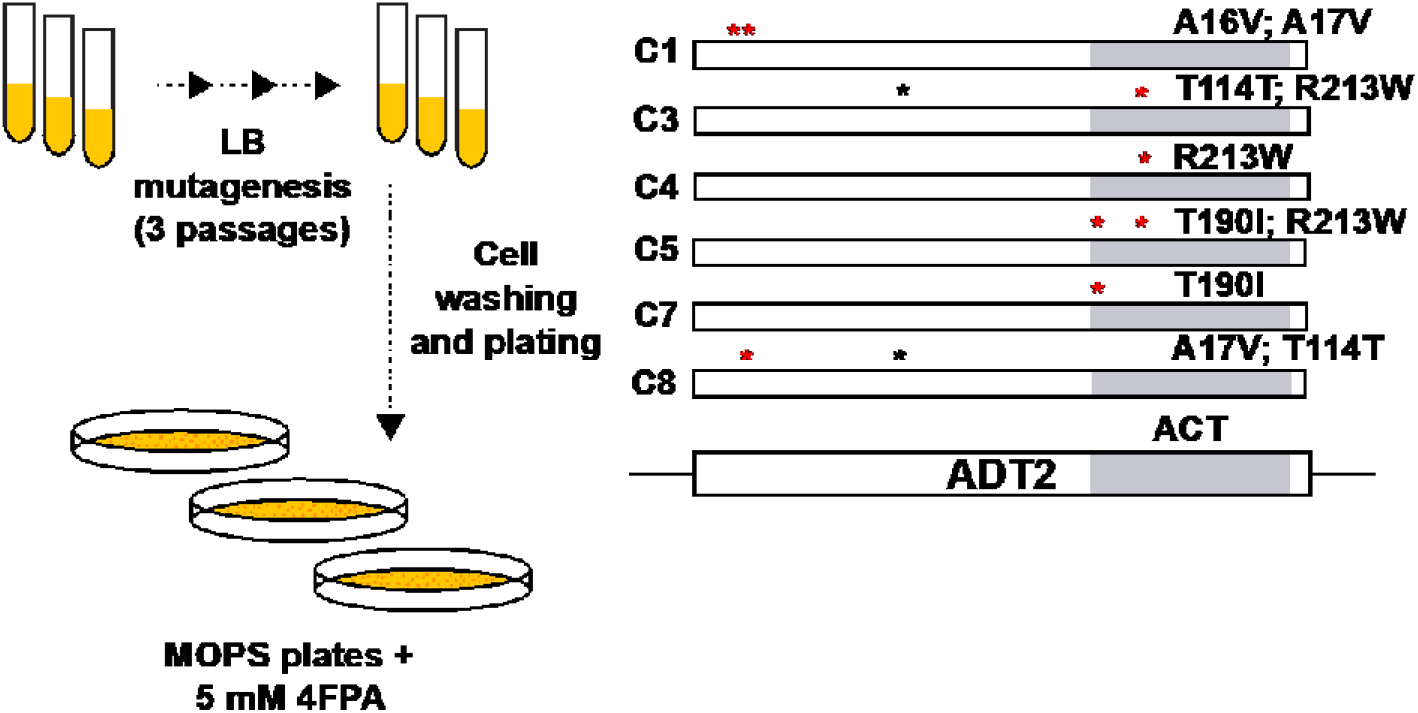
Trial of eMutaT7 in discontinuous mode. After accumulating a library of mutants in non-selective conditions in rich medium, cells were plated on selective minimal medium and the AtADT2 gene from resistant colonies was sequenced. Red asterisks are nonsynonymous mutations. Black asterisks are synonymous mutations. The ACT domain is shaded gray.

### eMuta in CDE mode

We next ran eMutaT7 in continuous mode and under selection,^4,12^ starting with ten independent populations and raising the 4FPA concentration stepwise from 0.5 to 30 mM during a total of 26 passages (**Figure 4A**). Each concentration step took five passages, during which growth of most populations improved (**Figure 4B**). Populations that showed no growth were replaced by dividing populations that did grow or, for selection steps up to 1 mM 4FPA, by populations maintained on minimal medium. Populations were plated as above on MOPS minimal medium plus 5 mM 4FPA at the 5 and 10 mM 4FPA steps during the campaign and when it ended at 30 mM. Bulk sequencing of cells from these plates showed three patterns in AtADT2 ORF mutations (**Table S4**).

**Figure 4.**
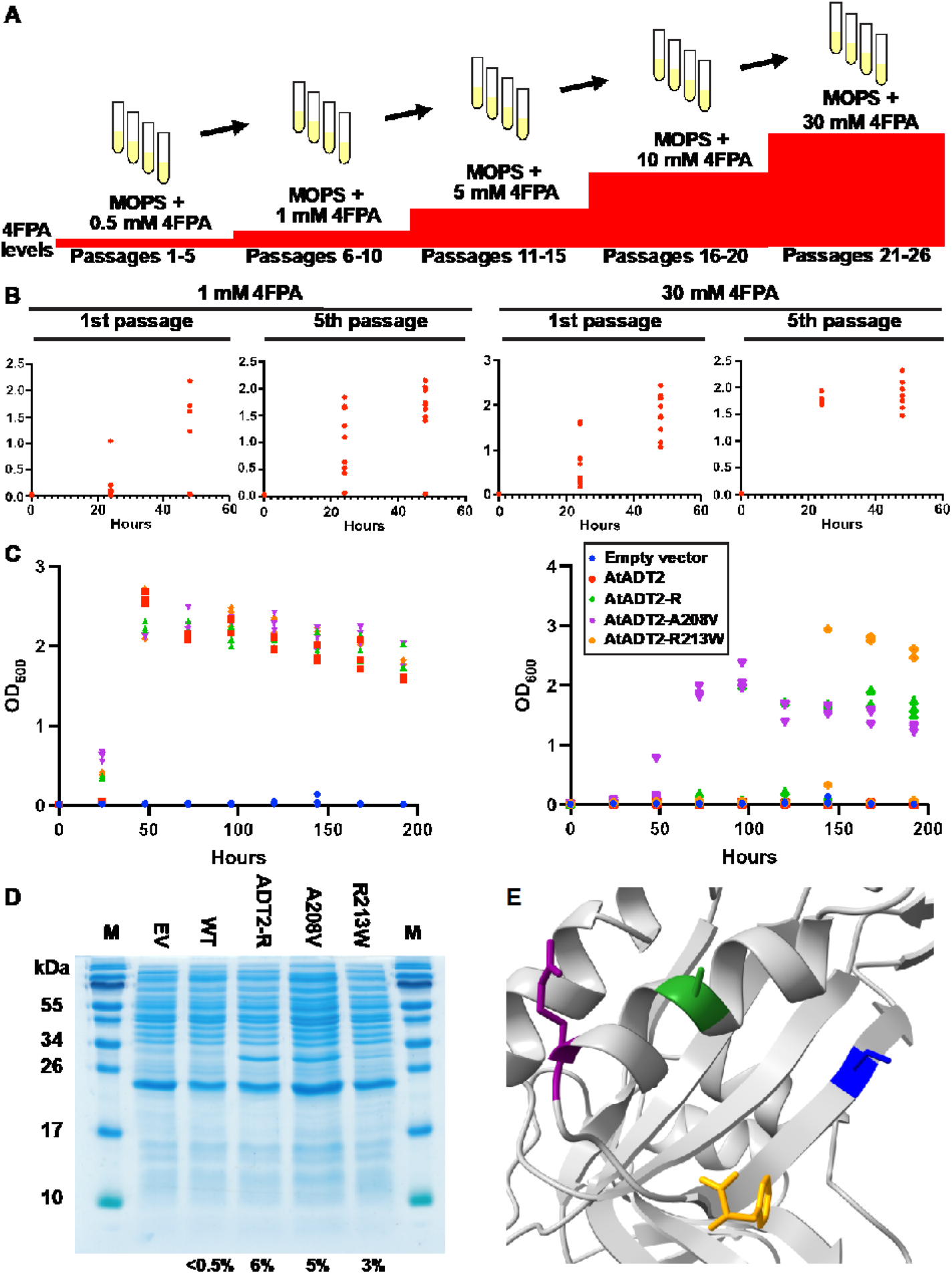
Operation of eMutaT7 in continuous mode and characterization of the mutants obtained. (A) Evolutionary campaign plan. 4FPA concentration was raised in five steps from 0.5 mM to 30 mM, with five passages at each step. (B) Data from the second (1 mM 4FPA) and final (30 mM 4FPA) steps illustrating how growth of most populations improves between first and last passages at these steps. Each datapoint is a separate population. (C) Expression levels of wildtype AtADT2 and of three variants; AtADT2 protein levels (percent of soluble protein) are given under the gel. (D) Growth in MOPS minimal medium alone or plus 5 mM 4FPA of *E. coli* strains expressing AtADT2 variants. Data are means ± s.e.m. of three independent replicates. EV, empty vector. (E) Structure model of the AtADT2 phenylalanine-binding pocket of the ACT domain showing positions of the benchmark S222N (blue), A208V (green), and R213W (violet) mutations that confer insensitivity to 4FPA.

First, A208V or R213W mutations, both in the ACT domain, had swept two or more independent populations by 5 mM 4FPA and persisted until the 30 mM step. Selection of these mutations in independent populations, and of R213W also in non-continuous mode (**Figure 3 and Table S3**), points to convergent evolution of fitness-enhancing mutations. Also pointing to convergent evolution, A208V and R213W both replaced other mutations over time. Second, one population (C6) was dominated at the 5 and 10 mM steps by a G201* nonsense mutation that truncates the ORF upstream of the regulatory ACT domain. As a G→T transversion, this mutation was probably not made by eMutaT7,^4,12^ i.e., was spontaneous. By the 30 mM step, the nonsense mutation had been displaced by a double mutant with the above A208V ACT domain mutation plus D240N, also in the ACT domain. The displacement suggests that the A208V plus D240N substitutions confer greater 4FPA resistance than the truncation. Third, while single mutants dominated all populations at the 5 mM step, by the 10 mM step one population was dominated by a double mutant, and there were three double mutants by the 30 mM step. Further, one of the mutations added between the 10 and 30 mM steps, A17V, was among those recovered in non-continuous mode (**Figure 3 and Table S3**). These patterns fit with sustained, target-specific hypermutagenesis by eMutaT7 throughout ~150 generations (26 passages × ~6 generations per passage). Sequencing the pHyo094 mutator plasmid at the end of the campaign confirmed the integrity of the T7RNAP-CD fusion, showing that mutations did not degrade or inactivate the mutational mechanism, despite the potential for this to happen.^8^

To validate the evolutionary campaigns, we compared the growth of strains expressing variants A208V or R213W with a strain expressing the known feedback-resistant mutant that was isolated from a mutagenized *Arabidopsis* population and carries the S222N mutation^12^ (first wrongly annotated as S222A^12^). We used a challenge of 5 mM 4FPA, which blocks growth of the strain expressing wildtype AtADT2 (**Figure 2A**). On minimal medium alone, the three strains expressing variants grew similarly to the strain expressing wildtype AtADT2, but adding 5 mM 4FPA led to marked differences (**Figure 4C**): relative to the benchmark S222N variant clones, those with the R213W variant grew similarly and those with the A208V variant grew faster. As expected, clones with the wildtype enzyme and clones harboring empty vector did not grow. Variation in growth between replicate clones is expected because their *AtADT2* genes are in the eMutaT7 system, and thus subject to ongoing hypermutation and hence to reduction or loss of AtADT2 expression.^8^ The S222N benchmark variant was more highly expressed in soluble form than wildtype AtADT2, as were the A208V and R213W variants to a lesser extent (**Figure 4D**). This is unlikely to relate to the observed resistance phenotypes because overexpression of feedback-inhibited enzymes cannot *per se* confer feedback-resistance,^18–20^ and the benchmark S222N variant gives feedback-resistance *in planta* without being overexpressed.^12^ Structure modeling based on the *Chlorobaculum tepidum* dehydratase (PDB code: 2QMX) suggested that, like the S222N mutation, the A208V and R213W mutations are located in the ACT domain ligand-binding pocket (**Figure 4E**). Based on the model, V208, W213, and N222 are all 11–12 Å from the phenylalanine ligand. The A208V and R213W mutations are thus positioned to disrupt phenylalanine binding, like S222N.

## CONCLUSIONS

CDE experiments require careful set-up, checks that the selection system is fit for purpose, and calibration of the dynamic range of selection.^4^ CDE of metabolic enzymes in *E. coli* additionally often requires the CDE system to work in minimal medium (which not all do^5^) and engineering of the platform strain.^21^ This study went through these steps for a realistic case of a plant enzyme and so provides a heuristic for future work of a similar kind. We established the following crucial points. First, eMutaT7 – and, by extension, probably other MutaT7 systems^4^ – can be used successfully in minimal medium. Second, pre-validation of the selection scheme with an enzyme having the desired property (i.e., a facsimile of the variants to be sought) is well worth the effort because it defines appropriate selection conditions at the outset. The same applies to checks for the likely frequency of cheaters. Third, bulk sequencing of the target gene in whole populations, which is fast and cheap, gives interim information on a campaign’s progress, and helps decide when to halt it, even if selected mutations are not yet fixed. Similarly, bulk sequencing of the mutator plasmid can detect failure, or incipient failure, of the mutational machinery and thereby dictate termination of a campaign instead of its futile prolongation.

The next major step in validating the directed evolution–genome editing (DE–GE)^9^ pathway to plant metabolic engineering is to return CDE-evolved AtADT2 variants such as A208V and R213W to the original plant, *Arabidopsis*. This will establish whether the variants confer the predicted phenotype (free phenylalanine accumulation^12^) and, more generally, how well they function in the plant host. As said at the outset, the differences between conditions in plant and prokaryote cells mean that it cannot be taken for granted that enzymes evolved in *E. coli* will necessarily function efficiently in a plant.

## METHODS

### Media and conditions

MOPS minimal medium (0.2% glycerol unless otherwise specified) was as described^22^ with added micronutrients.^23^ Antibiotics (µg mL^−1^) were: Carbenicillin (Carb) 100; Chloramphenicol (Cmp) 33; Kanamycin (Kan) 50; Apramycin (Apra) 50. Cells were washed in MOPS medium minus MgSO4, FeSO4, or micronutrients (basal MOPS). Growth was at 37°C, shaking at 250 rpm.

### Plasmid constructs

*AtADT2* and prephenate aminotransferase (*AtPAT*) were PCR-amplified from *Arabidopsis* cDNA generated using Superscript III (Invitrogen, Waltham, MA) and TA-cloned into pGEM-T Easy (Promega, Madison, WI). Both sequences were truncated to remove the plastid targeting peptide (**Table S5**); AtADT2 residue numbers in the text are for the truncated protein. The eMutaT7 target (pHyo182) and mutator (pHyo094) vectors were from Park and Kim.^11^ The *AtADT2* sequence and pHyo182 backbone were PCR-amplified and Gibson-assembled (NEB, Ipswich, MA). The S222N mutant of *AtADT2*^12^ was recreated by site-directed mutagenesis (Q5® Kit, NEB, Ipswich, MA). The empty vector was created by PCR-amplifying the pHyo182 backbone outside the start and stop codons of the AtADT2 ORF and ligating the ends. The cytidine deaminase domain was deleted from the control pHyo094 vector by site-directed mutagenesis as above. All constructs were sequence-verified.

### Strain engineering

The uracil-DNA glycosylase *ung* and *pheA* genes of *E. coli* MG1655 were deleted by recombineering;^24^ the Kan cassette from the corresponding Keio collection mutant was exchanged for the target gene, then removed via recombination of FRT sites. An expression cassette containing the pJ23101 promoter and B0032 ribosome binding site (iGEM collection) and the *rrnB* T1 terminator was cloned into the pGETS*flu*URA3 backbone^25^ via Gibson assembly. The *AtPAT* gene was then inserted into this cassette by Gibson assembly. The resulting expression cassette was PCR-amplified and recombineered into the *flu* locus as above, replacing Kan with Apra.

### *pheS* Mutagenesis in minimal medium

Single colonies of Δ*ung* cells harboring pHyo182-PheS and pHyo094 were inoculated in 3 mL of MOPS medium containing Carb and Cmp. Overnight cultures were diluted 100-fold into the same medium plus 0.2% arabinose and 0.1 mM IPTG. These cultures were grown for ~14 h and subcultured twice more. Cells were pelleted (4,000 g, 5 min), resuspended in 200 µL basal MOPS, diluted to OD^600^ 0.2, serially diluted ten-fold, and spotted (5-µL aliquots) on MOPS plates (Carb, Cmp, 0.2% arabinose, 0.1 mM IPTG, ± 10 mM *p*-chlorophenylalanine) and incubated overnight. Single colonies were restreaked on LB (Carb and Cmp) plates and grown overnight for sequencing of target gene amplicons.

### Complementation of the *ΔpheA* strain with AtPAT and AtADT2

Single colonies of the Δ*ung*::FRT Δ*pheA*::FRT Δ*flu*::AtPAT strain harboring pHyo094 and pHyo182 alone or containing *AtADT2* were inoculated into 3 mL MOPS medium (Carb, Cmp, 1 mM phenylalanine) and incubated overnight. Cells were pelleted as above, washed with 3 × 1 mL of basal MOPS, resuspended in 200 µL of basal MOPS, inoculated into 3 mL MOPS medium (Carb, Cmp, 0.002% arabinose, 1 mM IPTG) at an OD^600^ of 0.02, after which growth was monitored at OD^600^.

### 4FPA sensitivity calibration

Single colonies of the Δ*ung*::FRT Δ*pheA*::FRT Δ*flu*::AtPAT strain harboring pHyo094 and pHyo182 alone or containing *AtADT2* wild type or the feedback-resistant variant^12^ were inoculated into 3 mL MOPS medium (Carb, Cmp, 1 mM phenylalanine) and incubated overnight. Cells were pelleted, washed, resuspended, and inoculated into MOPS medium as above (complementation section) except that various concentrations of 4FPA were added. Growth was monitored as above.

### Cheater frequency test

Single colonies of the Δ*ung*::FRT Δ*pheA*::FRT Δ*flu*::AtPAT strain harboring pHyo182-AtADT2 and pHyo094 (T7RNAP-CD fusion or T7RNAP alone) were inoculated into 3 mL LB medium (Carb and Cmp) and incubated overnight. Cultures were diluted 100-fold into LB (Carb, Cmp, 0.1 mM IPTG, 0.2% arabinose), grown for 4 h, and subcultured twice more. Cells were pelleted, washed, resuspended, inoculated into MOPS, and grown as above. Cultures were then diluted to OD^600^ 1, and 100-µL aliquots were plated on MOPS medium (Carb, Cmp, 1 mM IPTG, 0.002% arabinose) containing 5 mM 4FPA; colonies were counted after four days. Aliquots were also diluted 10^5^-fold and plated on LB (Carb and Cmp) for total viable cell counts.

### Discontinuous mutagenesis

Procedures were as for cheater frequency tests except that the plasmids were pHyo182-AtADT2 and pHyo094. Colonies that grew in the presence of 5 mM 4FPA were restreaked on MOPS plates containing 5 mM 4FPA to minimize false positives; AtADT2 amplicons from resistant colonies were sequenced.

### Continuous mutagenesis with 4FPA selection

Ten single colonies of the Δ*ung*::FRT Δ*pheA*::FRT Δ*flu*::AtPAT strain harboring pHyo182-AtADT2 and pHyo094 were inoculated into 3 mL MOPS (Carb, Cmp, 1 mM phenylalanine), incubated overnight, washed, and resuspended as above. Cells were inoculated into 3 mL MOPS medium (Carb, Cmp, 1 mM IPTG, 0.002% arabinose) ± 0.5 mM 4FPA at starting OD^600^ 0.02. Cultures were grown for 48 h, monitoring OD^600^, and cells were pelleted, resuspended, and subcultured. 4FPA concentration was increased stepwise: 0.5, 1, 5, 10, and 30 mM (**Figure 4A**). Cultures that did not grow in 0.5 mM or 1 mM 4FPA were replaced with cells from paired MOPS cultures without 4FPA. Each 4FPA concentration was maintained for five passages. After passages 15, 20, and 26, cells were pelleted, washed, diluted, and plated as above on MOPS medium (Carb, Cmp, 1 mM IPTG, 0.002% arabinose) with 5 mM 4FPA; AtADT2 amplicons from resistant colonies were sequenced. AtADT2 mutants were tested for 4FPA resistance using the above growth protocol, inoculated into 5 mM 4FPA and grown for several days.

### DNA procedures

Target gene sequencing in clones (*pheS* mutagenesis and discontinuous *AtADT2* evolution) or populations (continuous *AtADT2* evolution) served as templates. Cells were resuspended in water and amplified by colony PCR using Phusion polymerase (Thermo Fisher Scientific, Waltham, MA). Mutant clones were restreaked on LB plates (Carb and Cmp) and mutant populations were maintained on MOPS plates (Carb, Cmp, 1 mM IPTG, 0.002% arabinose) containing 5 mM 4FPA. The T7 promoter plus *AtADT2* ORF were amplified using flanking primers and sequenced using internal primers. To sequence pHyo094, mutant populations were plated on LB (Carb and Cmp), then inoculated into LB (Carb and Cmp). Plasmids were isolated and sequenced using internal primers.

### Protein expression

Procedures were as in continuous mutagenesis except that cells were harvested at OD^600^ ~1, resuspended in 50 mM NaH^2^PO4, 300 mM NaCl, 5 mM DTT, pH 8.0 and sonicated (8 × 15-s pulse/45-s on ice). Lysates were cleared by centrifuging twice (22,000 g, 5 min). Protein was quantified by Bradford assay and 10 µg total protein analyzed by SDS-PAGE (15% acrylamide) and ImageJ software.

### Homology modelling

The truncated AtADT2 sequence was modeled against 2QMX structure using Phyre2. Models were aligned and analyzed with ChimeraX.

## Author Contributions

B.J.L. conceived the project with support from A.D.H. B.J.L. and A.D.H. designed experiments; B.J.L. performed the experiments. Both authors analyzed the data and wrote the manuscript.

## Notes

The authors declare no competing financial interest.

## Acknowledgments

This work was supported by NSF Fellowship IOS-2104739 to B.J.L., by USDA NIFA Hatch project FLA-HOS-005796, and by an Endowment from the C.V. Griffin, Sr. Foundation. We thank S. Kim for pHyo182 and pHyo094, and C.R. Reisch and L.Trujillo-Rodriguez for advice.

## REFERENCES

(1) Arnold, F. H. Directed evolution: bringing new chemistry to life. Angew. Chem. Int. Ed. Engl. 2018, 57, 4143–4148.

(2) Packer, M. S.; Liu, D. R. Methods for the directed evolution of proteins. Nat. Rev. Genet. 2015, 16, 379–394.

(3) Rix, G.; Liu, C. C. Systems for in vivo hypermutation: a quest for scale and depth in directed evolution. Curr. Opin. Chem. Biol. 2021, 64, 20–26.

(4) Molina, R. S.; Rix, G.; Mengiste, A. A.; Álvarez, B.; Seo, D.; Chen, H.; Hurtado, J.; Zhang, Q.; García-García, J. D.; Heins, Z. J.; Almhjell, P. J.; Arnold, F. H.; Khalil, A. S.; Hanson, A. D.; Dueber, vJ. E.; Schaffer, D. V.; Chen, F.; Kim, S.; Fernández, L. A.; Shoulders, M. D.; Liu, C. C. In vivo hyper-mutation and continuous evolution with cellular systems. Nat. Rev. 2022, 2, 1–22.

(5) García-García, J. D.; Joshi, J.; Patterson, J. A.; Trujillo-Rodriguez, L.; Reisch, C. R.; Javanpour, A. A.; Liu, C. C.; Hanson, A. D. Potential for applying continuous directed evolution to plant enzymes: an exploratory study. Life (Basel) 2020, 10, 179.

(6) Hendel, S. J.; Shoulders, M. D. Directed evolution in mammalian cells. Nat. Methods 2021, 18, 346–357.

(7) Rao, G. S.; Jiang, W.; Mahfouz, M. Synthetic directed evolution in plants: unlocking trait engineering and improvement, Synth. Biol. 2021, 6, ysab025

(8) García-García, J. D.; Van Gelder, K.; Joshi, J.; Bathe, U.; Leong, B. J.; Bruner, S. D.; Liu, C. C., Hanson, A. D. Using continuous directed evolution to improve enzymes for plant applications. Plant Physiol. 2022, 188, 971–983.

(9) Gionfriddo, M.; De Gara, L.; Loreto, F. Directed evolution of plant processes: towards a green (r)evolution? Trends Plant Sci. 2019, 24, 999–1007.

(10) Irfan, M.; Chavez, B.; Rizzo, P.; D’Auria, J. C.; Moghe, G. D. Evolution-aided engineering of plant specialized metabolism. aBIOTECH 2021, 2, 240–263.

(11) Park, H.; Kim, S. Gene-specific mutagenesis enables rapid continuous evolution of enzymes in vivo. Nucleic Acids Res 2021, 49, e32.

(12) Huang, T.; Tohge, T.; Lytovchenko, A.; Fernie, A. R.; Jander, G. Pleiotropic physiological consequences of feedback-insensitive phenylalanine biosynthesis in Arabidopsis thaliana. Plant J. 2010, 63, 823–835.

(13) Maeda, H.; Dudareva, N. The shikimate pathway and aromatic amino acid biosynthesis in plants. Annu. Rev. Plant. Biol. 2012, 63, 73–105.

(14) Lütke-Eversloh, T.; Stephanopoulos, G. Feedback inhibition of chorismate mutase/prephenate dehydrogenase (TyrA) of Escherichia coli: generation and characterization of tyrosine-insensitive mutants. Appl. Environ. Microbiol. 2005, 71, 7224–7228.

(15) Baas-Espinola, F. M.; Castro-Concha, L. A.; Vázquez-Flota, F. A.; Miranda-Ham, M. L. Capsaicin synthesis requires in situ phenylalanine and valine formation in in vitro maintained placentas from Capsicum chinense. Molecules 2016, 21, 799.

(16) Pine, M. J. Comparative physiological effects of incorporated amino acid analogs in Escherichia coli. Antimicrob. Agents Chemother. 1978, 13, 676–685.

(17) Liberles, J.S.; Thórólfsson, M.; Martínez, A. Allosteric mechanisms in ACT domain containing enzymes involved in amino acid metabolism. Amino Acids 2005, 28, 1–12.

(18) Chen, Q.; Man, C.; Li, D.; Tan, H.; Xie, Y.; Huang, J. Atrogenate dehydratase Isoforms differentially regulate anthocyanin biosynthesis in Arabidopsis thaliana. Mol. Plant 2016, 9, 1609–1619.

(19) Lütke-Eversloh, T.; Stephanopoulos, G. Combinatorial pathway analysis for improved L-tyrosine production in Escherichia coli: identification of enzymatic bottlenecks by systematic gene overexpression. Metab. Eng. 2008, 10, 69–77.

(20) Zhang, C.; Zhang, J.; Kang, Z.; Du, G.; Yu, X.; Wang, T.; Chen, J. Enhanced production of L-phenylalanine in Corynebacterium glutamicum due to the introduction of Escherichia coli wild-type gene AroH. J. Ind. Microbiol. Biotechnol. 2013, 40, 643–651.

(21) Bongaerts, N.; Edoo, Z.; Abukar, A. A.; Song, X.; Sosa-Carrillo, S.; Haggenmueller, S.; Savigny, J.; Gontier, S.; Lindner, A. B.; Wintermute, E. H. Low-cost anti-mycobacterial drug discovery using engineered E. coli. Nat. Commun. 2022, 13, 3905.

(22) Stewart, V.; Parales, J. Identification and expression of genes NarL and NarX of the Nar (nitrate reductase) locus in Escherichia coli K-12. J. Bacteriol. 1988, 170, 1589–1597.

(23) Neidhardt, F. C.; Bloch, P. L.; Smith, D. F. Culture medium for enterobacteria. J. Bacteriol. 1974, 119, 736–747.

(24) Datsenko, K. A.; Wanner, B. L. One-step inactivation of chromosomal genes in Escherichia coli K-12 using PCR products. Proc. Natl. Acad. Sci. USA 2000, 97, 6640–6645.

(25) Álvarez, B.; Mencía, M.; de Lorenzo, V.; Fernández, L. Á. In vivo diversification of target genomic sites using processive base deaminase fusions blocked by dCas9. Nat. Commun. 2020, 11, 6436.

