## Supplemental Figures and Tables for "Continuous directed evolution of a feedback-resistant *Arabidopsis* arogenate dehydratase in plantized *E. coli*": Figure S1.pdf

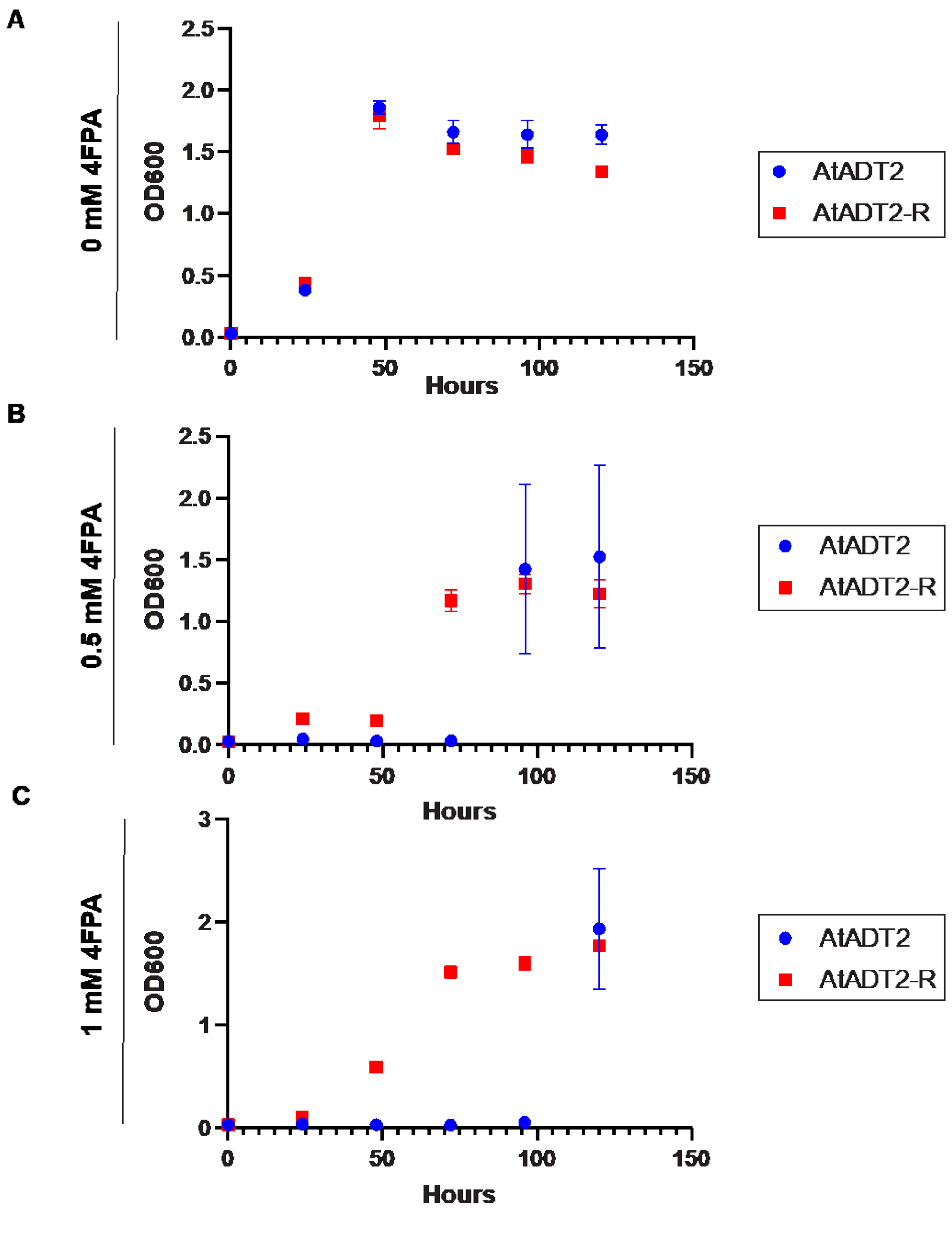


**Figure S1**. 4FPA-sensitivity of the plantized E. coli strain with wildtype AtADT2 and the feedback-insensitive variant AtADT2-R in (A) 0 mM, (B) 0.5 mM, and (C) 1 mM 4FPA. Data are means ± s.e.m. of three independent replicates.
